# The *Arabidopsis* outward K^+^ channel GORK mediates flg22-induced K^+^ efflux and stomatal closure

**DOI:** 10.64898/2026.10.10.758189

**Authors:** Limin Wang, Lili Yue, Ryan Toth, Sera Choi, Songyuan Zhang, Tobias Maierhofer, Simon Snoeck, Gijeong Kim, Kyle W. Bender, Yoonyoung Lee, Dietmar Geiger, Thomas A. DeFalco, Rainer Hedrich, Cyril Zipfel

**Affiliations:** Department of Plant and Microbial Biology, Zurich-Basel Plant Science Center, University of Zurich, Zurich, Switzerland; Molecular Plant Physiology and Biophysics, Julius-von-Sachs Institute, Biocenter, Julius-Maximilians-Universität Würzburg, Würzburg, Germany; The Sainsbury Laboratory, University of East Anglia, Norwich Research Park, Norwich, United Kingdom; Plant Breeding Institute, School of Life and Environmental Sciences, University of Sydney, Cobbitty, NSW, Australia; Department of Biology, Western University, London, ON, Canada; State Key Laboratory of Quantitative Synthetic Biology, Shenzhen Institute of Synthetic Biology, Shenzhen Institutes of Advanced Technology, Chinese Academy of Sciences, Shenzhen, China

**Author notes:** Correspondence and requests for materials should be addressed to C.Z. These authors contributed equally to this project (Limin Wang, Lili Yue).

## Abstract

Transient K^+^ efflux occurs rapidly following pathogen-associated molecular pattern (PAMP) perception by pattern recognition receptors (PRRs) as part of pattern-triggered immunity (PTI). However, the molecular identities of the K^+^ channels responsible for PTI-associated K^+^ efflux and the mechanisms linking immune signaling to K^+^ channel regulation remain largely unknown. Here, we show that the channel GUARD CELL OUTWARD RECTIFYING K^+^ (GORK) is required for K^+^ efflux and stomatal closure induced by flg22, a model PAMP derived from bacterial flagellin. BOTRYTIS-INDUCED KINASE 1 (BIK1), a receptor-like cytoplasmic kinase associated with immune receptor complexes that functions as a central regulator of PTI, interacts with GORK and phosphorylates the channel on the conserved T199 residue *in vitro.* Electrophysiological analyses indicate that BIK1 inhibits GORK activity in a kinase-dependent manner and that T199 plays an essential role for GORK function. Together, this study identifies GORK as the key K^+^ channel executing rapid PTI-associated K^+^ efflux and associated stomatal closure, thus shedding light on the mechanisms of early immune signaling.

## Introduction

Plants respond to biotic stresses by perceiving immune elicitors, such as pathogen-associated molecular patterns (PAMPs), through cell-surface pattern recognition receptors (PRRs), resulting in pattern-triggered immunity (PTI) (DeFalco and Zipfel, 2021; Ngou et al., 2022). Immediately downstream of elicitor perception is the rapid activation of receptor-like cytoplasmic kinases (RLCKs), most prominent among which is BOTRYTIS-INDUCED KINASE 1 (BIK1) (and the related kinase AvrPphB SUSCEPTIBLE 1-LIKE 1, PBL1), which functions as a central signaling hub by directly phosphorylating an array of downstream substrates, including the NADPH oxidase RESPIRATORY BURST OXIDASE HOMOLOG D (RBOHD), and multiple channels/transporters during PTI (Kadota et al., 2014; Li et al., 2014; Tian et al., 2019; Thor et al., 2020; Dindas et al., 2022; Yi et al., 2025). Recent analysis of BIK1 substrate specificity has further expanded its substrate repertoire, revealing diverse immune regulators involved in plasmodesmata function, intracellular nucleotide-binding leucine-rich repeat (NLR) receptor activation, and other defense-associated processes (Toth et al., 2026). These findings establish BIK1 as a master kinase that coordinates multiple layers of plant immune signaling.

Upon PTI activation, plants initiate a series of well-characterized immune responses, including reactive oxygen species (ROS) bursts, mitogen-activated protein kinase (MAPK) activation, immune gene expression and ion fluxes (DeFalco and Zipfel, 2021). While Ca^2+^ influx during PTI is being extensively studied; the role of K^+^ fluxes and the channels involved remain less understood. As the most abundant inorganic cation in plant cells, K^⁺^ plays essential roles in membrane potential regulation, turgor generation, and cellular metabolism (Shabala and Pottosin, 2014; Sardans and Peñuelas, 2021). Dynamic regulation of K^+^ homeostasis has emerged as an important component of plant immune responses. Transient K^+^ efflux is one of the earliest cellular responses following PAMP perception (Jeworutzki et al., 2010; Brauer et al., 2016), and recent spatial transcriptomic analyses further revealed that tissue-specific regulation of K^⁺^ transport is closely associated with immune responses (Wang et al., 2025). Despite these advances, the molecular identity of the K^⁺^ transporters responsible for PTI-associated K^⁺^ efflux and the mechanism by which immune signaling regulates K^⁺^ transports remain poorly understood.

The *Arabidopsis thaliana* (hereafter, Arabidopsis) genome encodes a family of Shaker K^⁺^ channels that are classified into five groups. The channels GUARD CELL OUTWARD RECTIFYING K^⁺^ CHANNEL (GORK) and the closely related homolog STELAR K^⁺^ OUTWARD RECTIFIER (SKOR) belongs to group V, and are the two major outward-rectifying K^⁺^ channels that mediate K^⁺^ efflux across the plasma membrane (Ache et al., 2000; Hosy et al., 2003; Demidchik et al., 2010; Lefoulon et al., 2016; Corratgé-Faillie et al., 2017; Förster et al., 2019). GORK is expressed in multiple tissues, including guard cells, roots, and vascular tissues, and plays a predominant role in mediating K⁺ efflux required for stomatal closure, and its activity is dynamically regulated by membrane depolarization, extracellular K⁺ concentration, ROS, and protein phosphorylation (Ache et al., 2000; Becker et al., 2003; Hosy et al., 2003; Demidchik et al., 2010; Lefoulon et al., 2016). In comparison, SKOR is thought to mediate primarily K^⁺^ loading into the xylem for long-distance K⁺ transport (Pilot et al., 1998; Hmidi et al., 2025). Previous electrophysiological study has shown that PAMP perception alters outward-rectifying K^⁺^ currents, suggesting that regulation of K^⁺^ channels is an integral component of stomatal immune responses (Zhang et al., 2008); however, whether GORK or SKOR contributes to PAMP-triggered K^+^ efflux and how its activity is regulated during immune activation remain unclear.

## Results

### In search of molecular identities responsible for K^+^ efflux upon flg22

The bacterial flagellin-derived peptide flg22 triggers K^+^ efflux in the mesophyll cells of Arabidopsis Col-0 (Jeworutzki et al., 2010); however, the molecular components responsible for this response remain elusive. Since the co-receptor BRI1-ASSOCIATED RECEPTOR KINASE 1 (BAK1) and downstream RLCKs BIK1/PBL1 are central signaling components immediately downstream of the flg22-FLAGELLIN SENSING 2 (FLS2)/BAK1 receptor complex (DeFalco and Zipfel, 2021), we tested whether they are required for flg22-induced K^⁺^ efflux. We measured net K^+^ fluxes in Arabidopsis mesophyll cells using non-invasive micro-test technology (NMT) following flg22 treatment. In wild-type (WT) Col-0, flg22 treatment triggered a significant K^+^ efflux, which was markedly reduced in the *bak1-5* (Schwessinger et al., 2011) and *bik1-10 pbl1-10* (Song et al., 2026) mutants (Fig. S1a, b). We further explored whether flg22-induced K^+^ efflux depends on outward rectifying K^+^ channels by investigating mutants *gork-2, skor-2* and *gork-2 skor-2* (generated by crossing *gork-2* and *skor-2*) (Fig. S1c-d) using the same technique. Compared with WT Col-0, both single mutants showed significantly reduced flg22-induced K^+^ efflux in mesophyll cell, while the double mutant exhibited further reduction (Fig. 1a, b), suggesting that GORK and SKOR cooperate to mediate flg22-induced K^+^ efflux. Since GORK plays an important role in guard cell K^⁺^ efflux and stomatal regulation (Ache et al., 2000; Hosy et al., 2003), we also tested the K^+^ fluxes in this cell type. flg22-induced K^+^ efflux was significantly reduced in *gork-2* as well as in two other CRISPR-Cas9 generated mutant lines (*gork-1c* and *gork-2c*) in guard cells (Fig. S2). Together, these results demonstrate that the activated FLS2 signaling complex (FLS2-BAK1-BIK1) as well as GORK and SKOR are required for flg22-induced K^⁺^ efflux. To determine whether GORK functions more broadly in early PTI signaling, we examined additional early signaling responses induced by flg22, such as Ca^2+^ influx and MAPK activation, but found that they were unaffected in the *gork-2* mutant (Fig. S3), suggesting that these signaling outputs work upstream or independently of K^+^ efflux.

**Figure 1.**
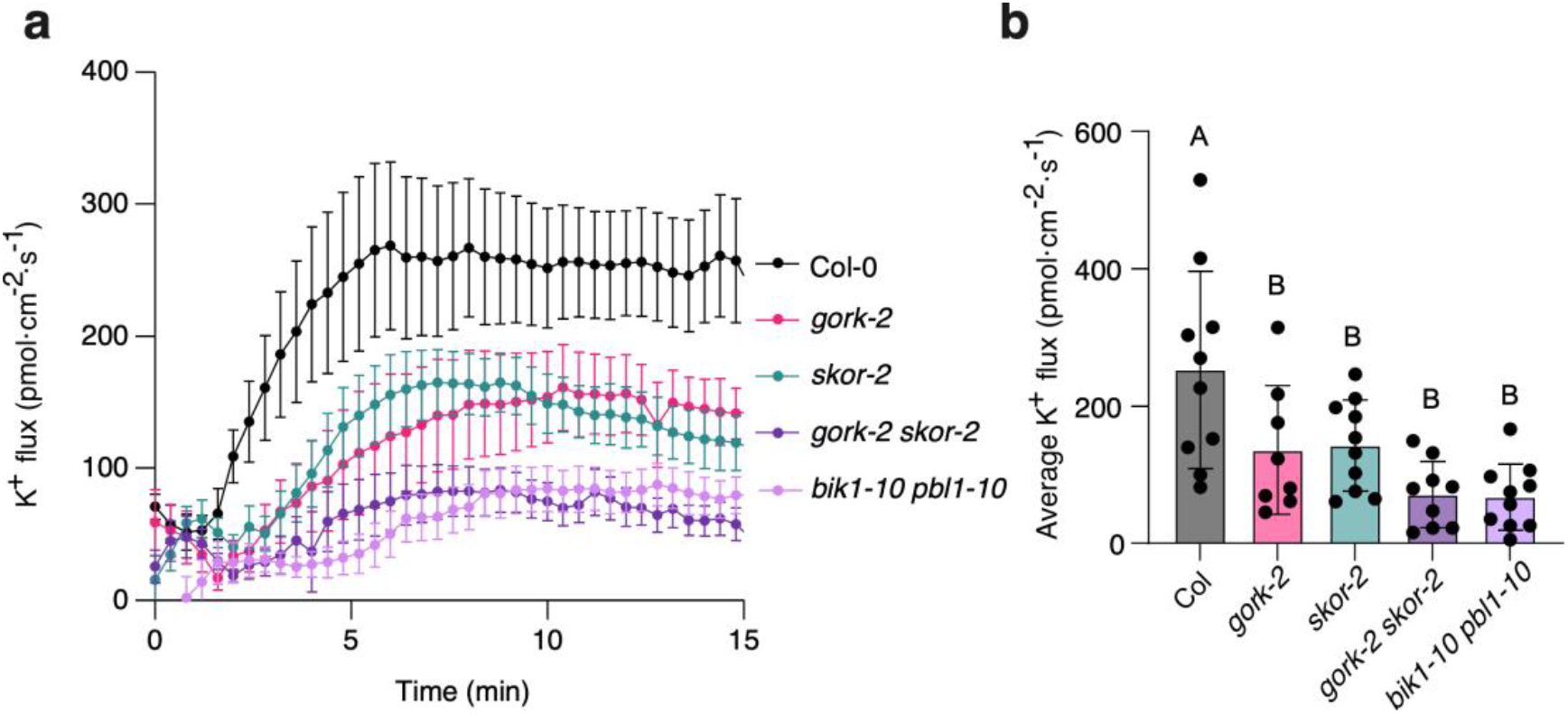
GORK and SKOR are required for flg22-induced K^+^ efflux. **a)** Time-course recordings of K^+^ flux in Arabidopsis mesophyll cells using NMT. Negative values indicate influx, and positive values indicate efflux. The flux recording started after 1 min treatment with 100 nM flg22. **b)** Quantitative analysis of average K^+^ fluxes from (a). Experiments were repeated 3 times with 8-10 individual leaves from each genotype analyzed. All data are presented as mean ± SE. Statistical significance was determined using one-way ANOVA with multiple comparisons.

### BIK1 interacts with GORK and trans-phosphorylates GORK at T199

Given the dependency on BIK1/PBL1 for flg22-induced K^+^ efflux (Fig. 1) and that BIK1/PBL1 directly phosphorylate ion channel/transporter proteins to regulate ionic conductivity during early PTI (Tian et al., 2019; Thor et al., 2020; Dindas et al., 2022; Yi et al., 2025), we tested the interaction between GORK/SKOR and BIK1. Using a split-ubiquitin yeast two-hybrid assay, we detected a physical interaction between BIK1 and GORK, but not between BIK1 and SKOR or ARABIDOPSIS THALIANA K^+^ RECTIFYING CHANNEL 1 (AtKC1), another member of the Shaker-type potassium channel family (Hedrich, 2012) (Fig. 2a; S4a, b). 5-FLUOROURACIL RESISTANCE 4 (FUR4), a yeast plasma membrane-localized multi-pass transmembrane protein, was included as a negative control. Reciprocal split-ubiquitin assays further validated the interaction between GORK and BIK1 (Fig. 2b). We further confirmed the interaction between BIK1 and GORK using a split-luciferase assay in *Nicotiana benthamiana* (hereafter, *N. benthamiana*) and co-immunoprecipitation (co-IP) assays in Arabidopsis protoplasts, respectively (Fig. 2c, d).

**Figure 2.**
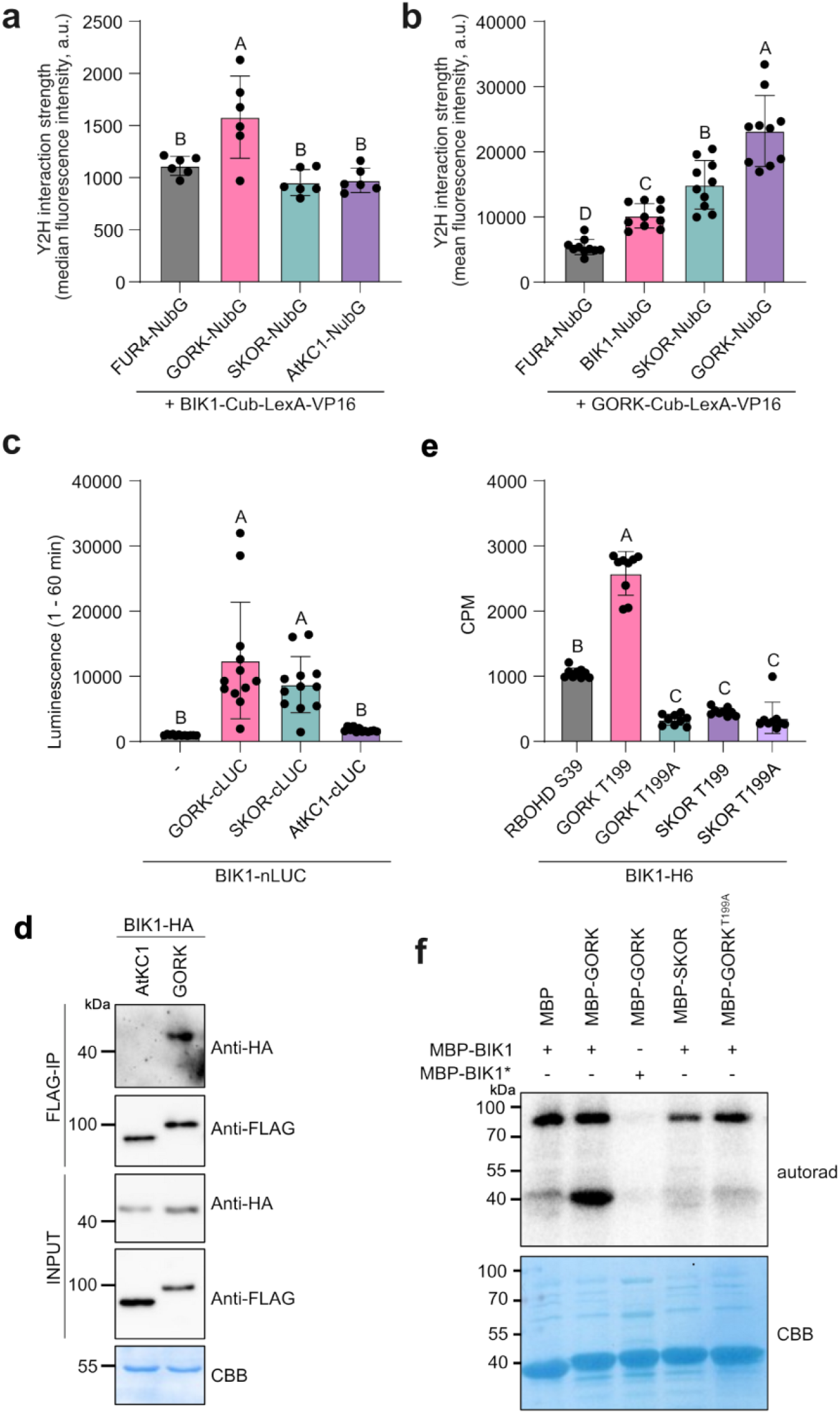
BIK1 associates with GORK and phosphorylates GORK at T199. **a-b)** Split-ubiquitin yeast two-hybrid assay between indicated proteins. FUR4 was used as negative control. **a)** Yeast cells co-expressing BIK1-Cub-LexA-VP16 together with the indicated NubG-fused membrane proteins were subjected to flow cytometry analysis. n = 6, error bars represent ± SE. **b)** Yeast cells co-expressing GORK-Cub-LexA-VP16 together with the indicated NubG-fused proteins were subjected to flow cytometry analysis. n = 10, error bars represent ± SE. **c)** Split-luciferase assay in *N. benthamiana*. Luminescence signals were continuously recorded from 1 to 60 min. Individual biological replicates are displayed as scatter dots. Proteins expression level in *N. benthamiana* were confirmed (Fig. S4c); n = 12. Error bars indicate ± SD. **d)** Co-IP assay between BIK1 and GORK in Arabidopsis mesophyll protoplasts. Total protein extracts (Input) were subjected to immunoprecipitation using anti-FLAG resin. Immunoblots were detected with anti-FLAG and anti-HA antibodies, respectively. Coomassie Brilliant Blue (CBB) staining serves as the loading control. **e)** *In vitro* peptide kinase assay with RBOHD-Ser39 WT, GORK-T199 WT, SKOR-T199 WT or mutant peptide substrates; n = 9. Error bars indicate ± SD. For a-d statistical significance was determined using one-way ANOVA with multiple comparisons. Different letters indicate statistical differences. **f)** *In vitro* radioactive kinase assays with the indicated combinations of purified recombinant proteins. Asterisk indicates kinase-dead variant. MBP serves as a negative control. All assays were repeated at least twice with similar results.

BIK1 is a highly selective kinase with preference to target Ser/Thr residues within a motif with a +3 Leu residue (DeFalco et al., 2022; Toth et al., 2026). Notably, GORK contains a S/T-X-X-L motif, T^199^-R-I-L, located in the cytoplasmic linker between the S4 and S5 domains (Fig. S5a, b), which is highly conserved amongst GORK homologs within the Brassicaceae family (Fig. S5c). We tested BIK1 trans-phosphorylation *in vitro* on peptide substrates derived from the conserved Thr in both GORK and SKOR and used a peptide substrate derived from known BIK1 target RBOHD’s S39 as a positive control (Kadota et al., 2014). BIK1 phosphorylated WT GORK, but not the phospho-ablative GORK^T199A^ mutant. In contrast, SKOR, which has +3 Val instead of Leu, lacked clear trans-phosphorylation by BIK1 (Fig. 2e). Corroborating the peptide kinase assay results, BIK1, but not a kinase-dead variant (K105A/K106A, hereafter, BIK1*), was able to phosphorylate the recombinant GORK cytosolic loop, whereas the phospho-ablative GORK^T199A^ and SKOR recombinant proteins showed no detectable phosphorylation (Fig. 2f). These *in vitro* data suggest that GORK is a BIK1 substrate and that T199 is a major phosphosite.

### BIK1 inhibits GORK channel activity in *Xenopus* oocytes

Having established GORK as a direct substrate of BIK1, we next asked whether BIK1 phosphorylation affects GORK channel activity. To test this, we expressed GORK alone or together with either WT BIK1 or the kinase-dead BIK1* and compared macroscopic K^+^ currents provoked by voltage stimulation. Expression of BIK1 or BIK1* alone did not produce detectable outward K^+^ currents (Fig. 3a). GORK expression generated a typical slow activating outward-rectifying K^+^ currents. Co-expression of WT BIK1 almost completely abolished GORK-mediated outward currents across the tested voltage range, whereas co-expression with the kinase-dead BIK1* had no detectable effect (Fig. 3a, b). Quantification of the steady-state current at +60 mV further confirmed that GORK currents were dramatically reduced by WT BIK1 but remained comparable to GORK alone in the presence of BIK1* (Fig. 3c), suggesting BIK1 inhibits GORK channel activity in a kinase-dependent manner.

**Figure 3.**
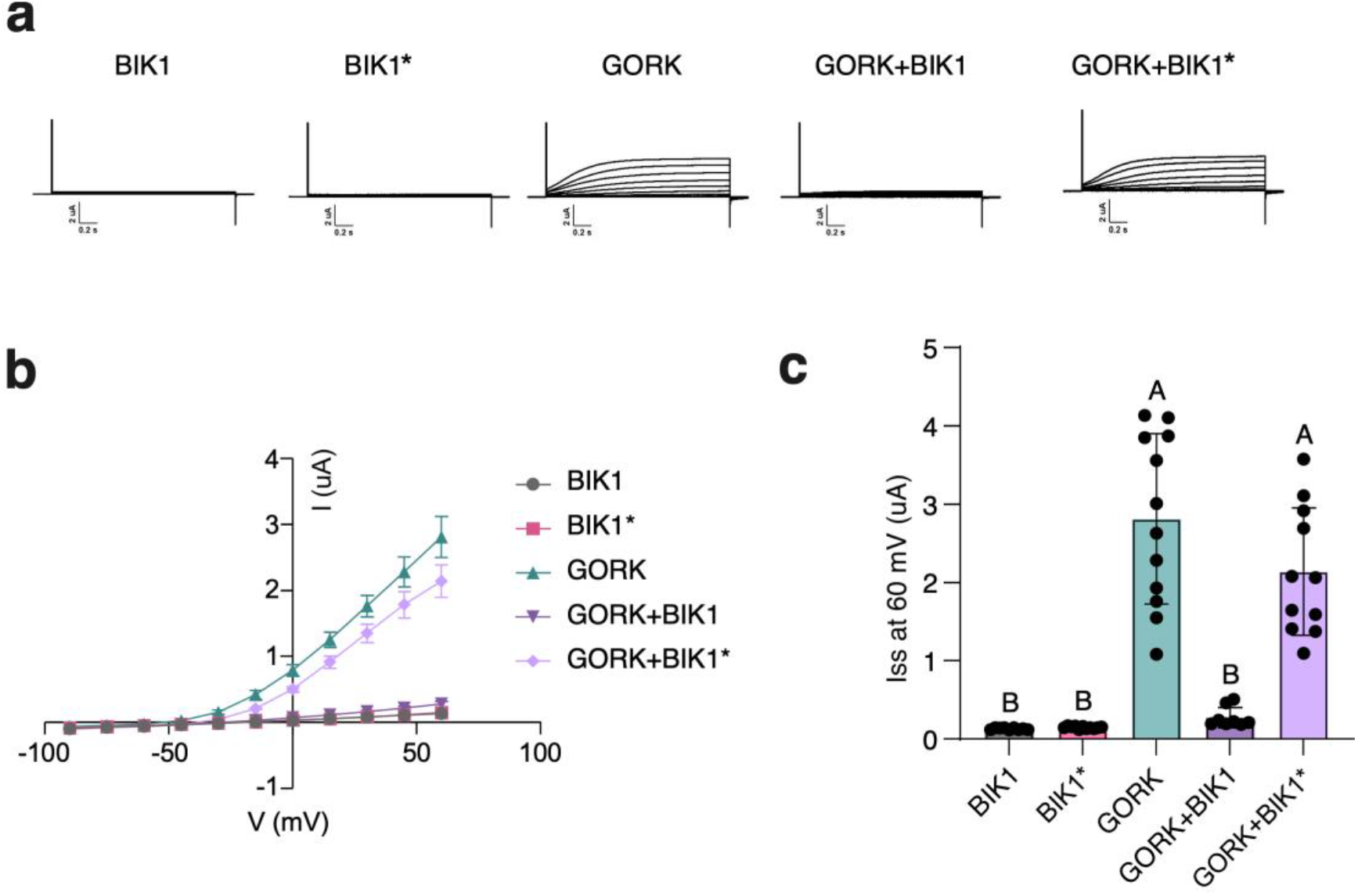
BIK1 inhibits GORK outward rectifying currents. **a)** Representative whole-cell current traces of oocytes injected with BIK1, BIK1*, GORK, or GORK co-injected with BIK1 or BIK1*. Voltage pulses were applied from - 90 to +60 mV with 15 mV increase; holding potential at -90 mV. **b)** Mean of current-voltage (I-V) relationships at steady state; n = 9-12. Error bars indicate ± SE. **c)** Comparison of steady state currents at 60 mV voltage; n = 9-12. Error bars indicate ± SD. Experiments were repeated twice with similar results. Statistical significance was performed using one-way ANOVA with multiple comparisons. Different letters indicate statistical differences.

### T199 is an important residue for GORK channel activity

To examine the impact of T199 phosphorylation on GORK function, we generated phospho-ablative (T199A) and phospho-mimic (T199D) variants of GORK. Both GORK^T199A^ and GORK^T199D^ displayed plasma membrane localization comparable to that of WT GORK in *Xenopus* oocytes (Fig. 4a), indicating that the amino acid substitutions did not affect GORK subcellular localization at the plasma membrane. Whole-cell current recordings revealed that the phospho-ablative GORK^T199A^ remained functional but exhibited significantly reduced outward K^+^ currents compared with WT GORK, whereas the phospho-mimic GORK^T199D^ variant exhibited almost no detectable channel activity (Fig. 4a, b). Quantification of the steady-state current at +60 mV confirmed a reduction in channel activity of GORK^T199A^ and GORK^T199D^ (Fig. 4c). We then introduced WT GORK, GORK^T199A^ and GORK^T199D^ into the *gork-2* mutant background using native promoter (Fig. S6) and measured flg22-induced K^+^ efflux using NMT. WT GORK fully restored flg22-induced K^+^ efflux in the *gork-2* mutant, whereas GORK^T199A^ or GORK^T199D^ did not, suggesting that T199 is essential for GORK induced K^+^ efflux during flg22 signaling (Fig. 4d, e).

**Figure 4.**
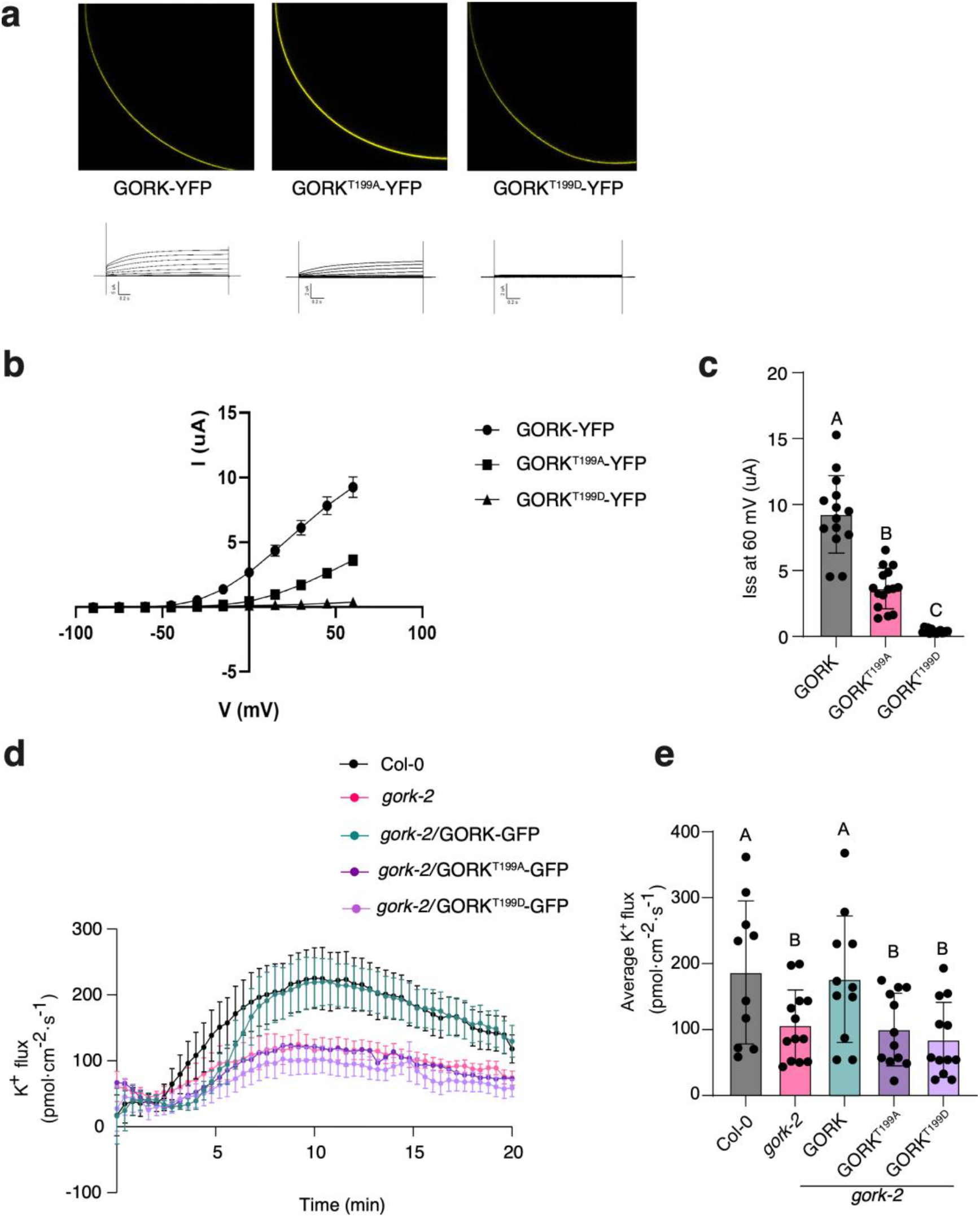
T199 is critical for GORK channel activity. **a)** Subcellular localization (upper) and representative whole-cell current traces (lower) of GORK-YFP, GORK^T199A^-YFP, and GORK^T199D^-YFP expressed in *Xenopus* oocytes. **b)** Mean of current-voltage (I-V) relationships at steady state; n = 14-15. Error bars indicate ± SE. **c)** Comparison of steady state currents at 60 mV voltage; n = 9-12. Error bars indicate ± SD. **d)** Time-course recordings of K^+^ flux in Arabidopsis mesophyll cells using NMT. The flux recording started after 1 min treatment with 100 nM flg22. **e)** Quantitative analysis of average K^+^ fluxes from (d). Experiments were repeated 3 times with 10-13 individual leaves from each genotype were analyzed. All data are presented as mean ± SE. Statistical significance was determined using one-way ANOVA with multiple comparisons. Different letters indicate statistical differences.

### GORK is required for flg22-induced stomatal closure

Stomatal closure represents one of the earliest defense responses during PTI, restricting pathogen entry (Melotto et al., 2006). Since guard cell K^+^ efflux is essential for reducing turgor pressure and driving stomatal closure (Jezek and Blatt, 2017), we next investigated whether GORK-mediated K^+^ efflux contributes to flg22-induced stomatal closure. Stomatal aperture was measured microscopically following flg22 treatment, with abscisic acid (ABA)-induced stomatal closure as a control to distinguish PTI-specific defects from general impairments in stomatal movement. In WT Col-0 leaves, both flg22 and ABA significantly reduced stomatal aperture (Fig. 5a, b). In *gork-2*, however, flg22-induced stomatal closure was abolished, whereas ABA still induced a significant reduction in stomatal aperture (Fig. 5a, b), suggesting that GORK is specifically required for flg22-triggered stomatal closure under these conditions. This phenotype was independently confirmed in two CRISPR-Cas9-generated *gork-1c* and *gork-2c*mutant lines (Fig. 5c). Complementation of *gork-2* with *pGORK::GORK-GFP*fully restored flg22 responsiveness, whereas expression of GORK^T199A^ or GORK^T199D^ variants did not (Fig. 5d). Collectively, these results demonstrate that GORK channel activity is required for flg22-induced stomatal closure and identify T199 as a critical residue linking GORK activation to stomatal closure in this context.

**Figure 5.**
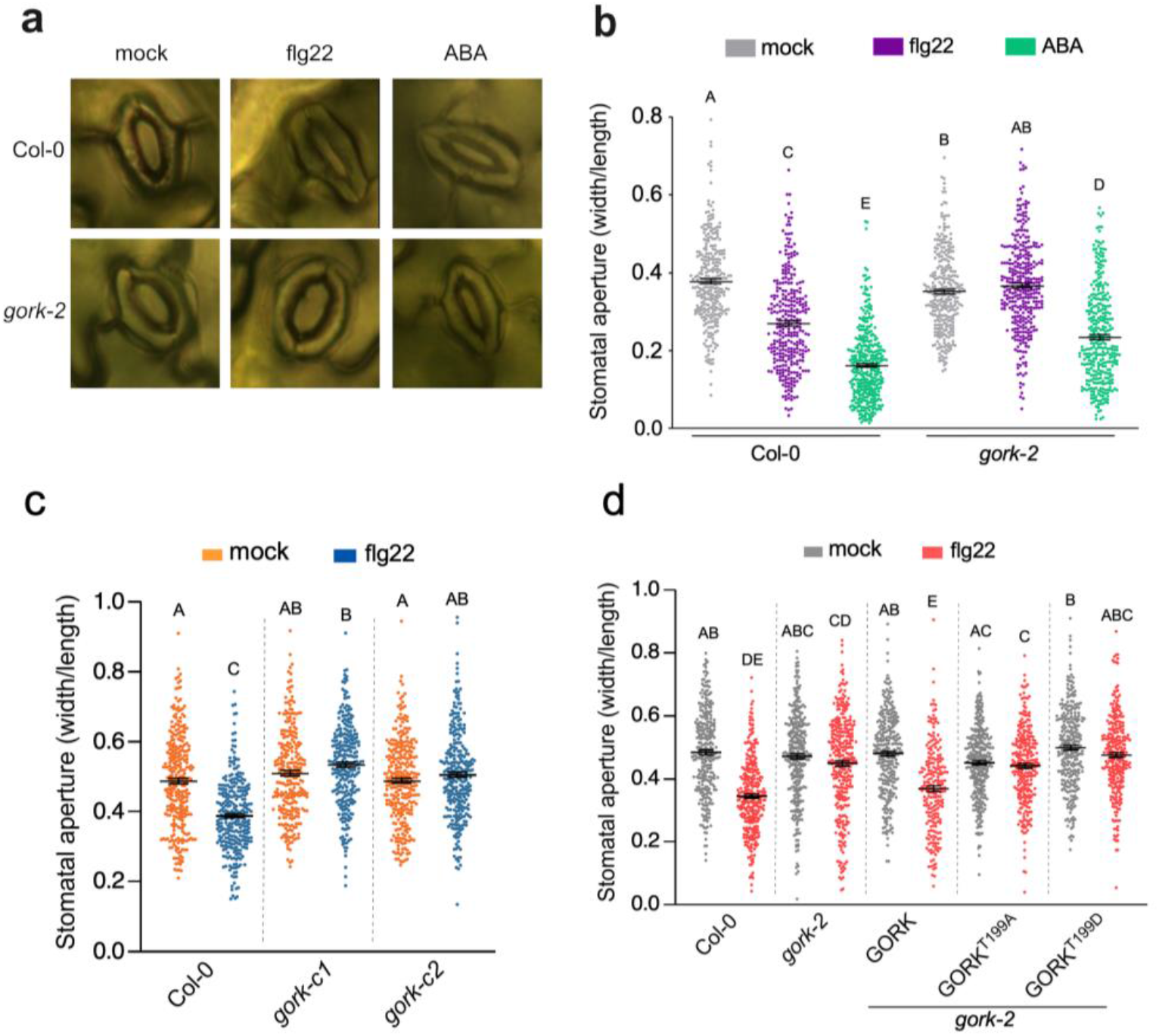
GORK is required for stomatal closure. **a)** Representative images of stomatal apertures in Col-0 and *gork-2* treated with 5 μM flg22, 10 μM ABA, or water for 2.5 h. **b)** Quantification of stomatal apertures shown in (a). Individual data points are shown as mean ± SD, from 4 individual plants with total 8 leaf discs sampled (n = 141-191 stomata). The experiment was performed twice with similar results. **c)** Stomatal aperture measurement of complemented lines. Experiments were repeated 3 times with 15 individual leaves from each genotype. All data are presented as mean ± SE. **d)** Stomatal aperture measurement of mutant lines *gork-1c* and *gork-2c*. Experiments were repeated three times with 11 individual leaves from each genotype. All data are presented as mean ± SE. Statistical significance in (b-c) were performed using one-way ANOVA with multiple comparisons.

## Discussion

In this study, we identified GORK as the main channel mediating flg22-induced K^+^ efflux and stomatal closure. We demonstrate that BIK1 directly associates with GORK and inhibits GORK channel activity in a kinase-dependent manner. Genetic complementation experiments further demonstrate that the conserved T199 residue that is phosphorylated by BIK1 *in vitro* is indispensable for GORK-dependent flg22-induced K^+^ efflux and stomatal closure.

Of note, in our conditions, using whole leaf discs, GORK was not required for ABA-induced stomatal closure, suggesting that different K^+^ channels are involved in immune (flg22) and ABA signaling in guard cells. This is similar to our previous observation that the BIK1-targeted Ca^2+^-permeable channels OSCA1.3 and OSCA1.7 are required for flg22-induced stomatal closure but dispensable for that induced by ABA (Thor et al., 2020). Importantly, while GORK was previously suggested to function in ABA signaling, this was based on stomatal closure assays using epidermal peels with a Ws-0 *gork-1* allele (Hosy et al., 2003), or seed germination and seedling growth assays with the Col-0 *gork-1* (SALK_082258) and *gork-2* (SALK_144737) alleles (Lim et al., 2019). Yet, mirroring our findings, another Col-0 mutant allele, *gork1-2* (GABI_865F05), was found to be still responsive to ABA, while being insensitive to jasmonic acid in stomatal closure assays (Förster et al., 2019). In this latter study, *gork1-2* stomata were however still able to close in response to flg22, while our results with *gork-2* (SALK_144737), newly generated *gork* CRISPR mutants (*gork-1c* and *gork-2c*) and complementation lines clearly show that GORK is required for flg22-induced stomatal closure. The availability of novel *gork* mutant alleles and complementation lines might allow reassessing the role of GORK in diverse pathways, similarly to as was recently did with BIK1 (Song et al., 2026).

Several characterized BIK1 substrates, including RBOHD and the calcium-permeable channels OSCA1.3 and CNGC2/3/4, are positively regulated by BIK1 phosphorylation to promote early immune signaling (Kadota et al., 2014; Tian et al., 2019; Thor et al., 2020; Yi et al., 2025). However, BIK1 was found to inhibit the function of several other substrates, including the primary phosphate transporter PHT1;4 (Dindas et al., 2022), protein phosphatases of the POLTERGEIST-LIKE family (DeFalco et al., 2022), and several sensor NLRs (Toth et al., 2026) . Our results showing that phosphorylation by BIK1 suppressed GORK-mediated outward K^+^ currents. At the same time, the *bik1-10 pbl1-10* mutant show reduced flg22-induced K^⁺^ efflux. A possible explanation is that BIK1 is required for the initiation of early immune responses, including cytosolic Ca²⁺ elevation, depolarization and ROS production which are expected to contribute to the membrane depolarization required for GORK activation (Ache et al., 2000; Demidchik et al., 2003; Trouverie et al., 2008; Demidchik et al., 2010; Geiger et al., 2010; Demidchik, 2018; Pottosin et al., 2018). Once GORK is activated, phosphorylation by BIK1 may subsequently attenuate channel activity, thereby limiting excessive K^+^ loss. Such a dual regulatory mechanism would allow rapid activation of K^+^ efflux while preventing prolonged channel opening, ensuring tight control of ion homeostasis during immune signaling.

Our results identify T199 as a critical regulatory residue for GORK channel activity in *Xenopus* oocytes and Arabidopsis. The reduced channel activity of the both phospho-ablative and phospho-mimic mutant suggests that T199 itself is required for normal channel gating, consistent with its location within the S4–S5 cytoplasmic linker, a key mechanical coupling element that translates voltage-sensing domain (S4) movement into opening of the pore domain (S5-S6) in the Shaker-type K^⁺^ channels. Recent structural analyses of GORK further highlight the S4-S5 linker as a critical interface governing electromechanical coupling between voltage sensing and pore gating (Lu et al., 2002; Zhang et al., 2025). Structural predictions revealed that T199A likely affects the channel activity of GORK by disrupting the interaction between S4-S5 and the C-helix; T199D might tightly interact with R320 from the C-helix to force a local inactive conformation (Fig. S7). The dual role of T199 as both a structural determinant of channel gating and a potential target of BIK1 phosphorylation complicates the interpretation of mutational analyses, making it difficult to distinguish the structural contribution of this residue from the direct functional consequences of its phosphorylation. Whether additional BIK1-dependent phosphorylation sites contribute to GORK regulation remains an open question. Identifying such sites, if present, will help clarify how BIK1-mediated phosphorylation regulates GORK channel activity. It will also be interesting to investigate whether additional kinases also regulate GORK activity during immunity, as previously proposed during abiotic stress signaling (Corratgé-Faillie et al., 2017; van Kleeff et al., 2018; Förster et al., 2019).

The genetic functional overlap between GORK and SKOR has been observed under abiotic stress conditions, where both channels contribute to salt stress responses and root growth regulation (Hiya et al., 2024). Consistent with their potential functional overlap, our results reveal both channels contribute to flg22-induced K^+^ efflux; however, GORK and SKOR are differentially regulated, with only GORK being phosphorylated by BIK1. These findings identify GORK as a BIK1-associated K^+^ channel, while the potential contribution and regulation of SKOR remain to be further defined. Given previous study and our results (Fig. 2b) showing that SKOR and GORK physically interact and assemble into heteromeric K^⁺^ efflux channel complexes (Dreyer et al., 2004), and that heteromeric assembly is a common feature of plant shaker channels (Xicluna et al., 2007; Yuan et al., 2024), it will be important to determine whether such channel organization contributes to the regulation of K^+^ efflux during immune signaling.

## Methods

### Plants and growth conditions

The details of the plant materials used in this study are shown in Supplementary Table S1. *Arabidopsis thaliana* ecotype Columbia-0 (Col-0) served as the wild-type (WT) genetic background throughout this study.

For sterile culture, all seeds were subjected to surface sterilization with chlorine gas for 5 to 6 h. Sterilized seeds were plated on half-strength Murashige and Skoog (½ MS) medium, 1 % (w/v) sucrose, and 0.9 % (w/v) agar. Following stratification in darkness at 4 °C for 2-3 days, seeds were transferred to a growth chamber set to a 16-h light/8-h dark photoperiod, with corresponding temperatures of 22 °C (day) and 18 °C (night). For soil cultivation, 10-day-old seedlings were transplanted into potting soil and maintained in a controlled growth chamber under the following conditions: 150 µmol·m⁻²·s⁻¹ light intensity, 60 % relative humidity, 20 °C constant temperature, and a 10-h photoperiod. *N. benthamiana* plants were grown in a controlled environment chamber at 150 µmol light intensity, 45-60 % relative humidity and 25 °C in a 16-h light/8-h dark cycle.

### Plasmid construction and generation of transgenic plants

The details of the constructs and primers used in this study are shown in Supplementary Tables S2 and S3, respectively.

For CRISPR/Cas9 cloning, guide RNAs (gRNAs) targeting *GORK* (gRNA1c: CTCCATGGGAGTGAACAATGAGG; gRNA2c: AGAGTCTCTCAGGCAGACCGCGG) were designed using the CHOPCHOP (v3) tool (Labun et al., 2019) and assembled into the pAGM55261 vector via the Golden Gate cloning. For *Xenopus* oocytes expression, the target sequences were cloned into pGEMHE vector by restriction digestion and ligation. For stable transgenic line generation, coding and promoter sequences PCR-amplified from Arabidopsis genomic DNA were subcloned into p641-Esp3I. Golden gate assembly was performed in pICSL86955OD with pICH41421 (NOS terminator) and the suitable C-terminal tag module to assemble promoter-coding expression cassettes. The resulting binary vectors were transformed into Arabidopsis using *Agrobacterium tumefaciens* strain GV3101 via the floral dip method (Zhang et al., 2006). For Arabidopsis protoplast expression, the target sequences were subcloned into the p641 entry vector, followed by golden gate assembly with the pICSL50007 (C-terminal 3×FLAG tag) and pICSL50009 (C-terminal 6xHA tag) into the modified HBT promoter-driven protoplast expression vector (Toth et al., 2026). Constructs for *N. benthamiana* transient expression for split luciferase assays were generated using Golden Gate assembly into pICSL86955OD with pICH51277 (35S promoter), pICSL01002 (C-terminal N-Luc tag) and pICSL01003 (C-terminal C-Luc tag). Constructs for expression of recombinant protein in *E. coli* were generated by inserting PCR fragments into pOPINM using InFusion (Takara). The phospho-ablative GORK^T199A^ and phospho-mimic mutant GORK^T199D^ were generated by site-directed mutagenesis.

### Protein extraction and MAPK activation assay

The assay was performed as previously described (Zhai et al., 2024). Briefly, plant tissues were ground in liquid nitrogen and lysed with protein extraction buffer (50 mM Tris-HCl pH 7.5, 100 mM NaCl, 10 % glycerol, 2 mM EDTA, 5 mM dithiothreitol, 1 % IGEPAL, 2 mM sodium molybdate, 1 mM sodium fluoride, 1 mM sodium orthovanadate, 4 mM sodium tartrate, protease inhibitor cocktail). After centrifugation at 4 °C, supernatants were collected and separated by SDS-PAGE before transfer onto PVDF membranes. Membranes were blocked and incubated with corresponding primary and HRP-conjugated secondary antibodies; target proteins were detected via ECL chemiluminescence. Coomassie brilliant blue staining served as the loading control. All antibodies and consumables were as described previously (Zhai et al., 2024).

### Ion flux measurement using NMT

Non-invasive Micro-test Technology (NMT; YoungerUSA) coupled with imFluxes v.3.0 software (YoungerUSA) was used to measure net ion flux across the plasma membrane (Liu et al., 2023). Four-week-old plants were used for analysis. After the epidermis peeling, leaves were incubated in the solution (0.1 mM KCl, 0.1 mM CaCl_2_, 0.1 mM MES pH 6) at room temperature overnight before measurement. Prior to measurement, the solution was refreshed, and leaves were further incubated for an additional 30 min. Microelectrodes were pulled from GC150T-10 borosilicate glass capillaries (Harvard Apparatus), heated at 220 °C for 3 h and silanized with *N,N*-dimethyltrimethylsilylamine (Sigma, 41716). Electrodes with 1-2 μm tip apertures were filled with ionophore at the tip and backfilled with 100 mM KCl or CaCl_2_. After calibration against standard solutions, the microelectrode was fixed 20 μm above individual cell and scanned reciprocally between 20 μm and 60 μm.

### Injection of *Xenopus* oocytes and two-electrode voltage clamp (TEVC) measurements

The *Xenopus* oocyte preparation and injection were performed as previously described (Förster et al., 2019). In brief, oocytes were microinjected with cRNA encoding wild-type or mutant GORK/BIK1-GFP fusion proteins. All oocytes were cultured in ND96 medium at 16 °C for 2-3 days to permit protein translation prior to confocal imaging and electrophysiological detection. GORK channel activity was recorded in TEVC measurements on oocytes exposed to a bath solution (10 mM KCl, 90 mM LiCl, 1 mM CaCl_2_, 1 mM MgCl_2,_ 10 mM Hepes-NaOH pH 7.5). Electrophysiological data acquisition was performed with Patchmaster (software version 2×92; Multi Channel Systems MCS GmbH, Germany). Electrophysiological data were analyzed with the software Igor Pro 8 (waveMetrics, Inc., Lake Oswego, Oregon, USA).

### Stomatal aperture measurement

Four-week-old plants were used for analysis. Leaf discs were incubated in the opening buffer (10 mM KCl, 50 μM CaCl_2_, 10 mM MES-KOH pH 6.15) under light for 2 h with the lower surface facing up. Then, the leaf discs were treated with 5 μM flg22 for 2.5 h under the same light conditions. The opening buffer was removed, and the leaf discs were immediately immersed in the fixation solution (a mixture of 10 mL 4 % formaldehyde and 30 mL 1×PBS) for 1-2 min to halt guard cell movement. Gently rotate the petri dish to ensure the samples are fully immersed in the fixation solution. After two washes with PBS, mount the leaf discs on slides and cover them with coverslips. Stomatal apertures were photographed using a light microscope.

### Split luciferase assay

The assays were conducted following the previously reported protocol with slight modifications (Chen et al., 2008). Briefly, the designated vector constructs were transiently expressed by infiltration into leaves of 4- to 5-week-old *N. benthamiana* plants. At 2 day-post infiltration (2 dpi.), 4 mm leaf discs were excised and placed in 96-well plates containing double-distilled water. After a 6-h equilibration period, the water was replaced with 1 mM luciferin solution, and the plates were incubated in darkness for 1 h. Luminescence signals were subsequently quantified using a Tecan Spark microplate reader.

### Recombinant protein expression and purification

Unless specified otherwise, all recombinant proteins were expressed in *E. coli* strain BL21(DE3) Rosetta pLysS. Purification was carried out using PureCube Co-NTA agarose (Cube Biotech).

### *In vitro* kinase assays and autoradiography quantification

For *in vitro* kinase assays using recombinant proteins as substrate, approximately 1 µg of BIK1 and 2 µg substrate protein were mixed in kinase reaction buffer (25 mM Tris-Cl pH 7.4, 5 mM MnCl_2_, 5 mM MgCl_2_, 1 mM dithiothreitol). Reactions were initiated by addition of 5 µM ATP containing 0.5 µCi γ-^32^P-ATP, in a total reaction volume of 30 µL. Reactions were stopped following 30 min at 25 °C by addition of SDS-loading dye and heating at 70 °C for 10 min. Reactions were separated by SDS-PAGE, transferred to PVDF membrane, and stained with CBB G-250. Autoradiographs were imaged using an Amersham Typhoon Phosphorimager (GE Healthcare).

For *in vitro* kinase assays with peptide substrates, approximately 1 µg of BIK1 and 100 µM substrate peptides were mixed in kinase reaction buffer. Reactions were initiated upon addition of 100 µM ATP containing 0.1 µCi γ-^32^P-ATP, in a total reaction volume of 25 µL. Following 20 min at 25 °C, 20 µL of reaction was transferred to a 1-cm^2^ piece of cation exchange paper (SVI-P). Paper was washed 3 times in 0.5 % phosphoric acid. ^32^P signal was measured as counts per minute (cpm) on a scintillation counter.

### Quantitative split-ubiquitin membrane yeast two-hybrid assay

Split-ubiquitin yeast two-hybrid assay was performed as previously described (Kavitha Iyer et al., 2005), with custom modifications to enable quantitative measurement of protein-protein interaction strength. Briefly, a fluorescent reporter yeast strain was generated via genomic integration of plasmid FRP795 into the FRY70 parental strain, as previously described (Ottoz et al., 2014). GORK, SKOR, AtKC1, BIK1 with C-terminal Cub-LexA-VP16 and NubG fusions were used as baits and preys, respectively. Yeast membrane proteins FUR4 with C-terminal NubG fusions were used as negative control. At least six independent colonies of each bait-prey pair were cultured in yeast synthetic drop-out medium until exponential growth phase. Fluorescence was measured using flow cytometer BD FACSCanto™ II equipped with a High Throughput Sampler (HTS). Mean and median fluorescence intensity values of at least 10,000 single cells were calculated by BD FACSDiva™ software and used to quantify the interaction strength.

### Arabidopsis protoplast isolation, transfection and co-immunoprecipitation assay

This part was performed as previously described (Wu et al., 2009; Toth et al., 2026). Protoplasts were isolated from 5-week-old Arabidopsis rosette leaves. The leaf epidermal layers were stripped off with 3M™ adhesive tape, and the peeled leaf tissues were digested in enzyme solution containing 1.5 % cellulase R10, 0.4 % macerozyme, 0.4 M mannitol, 20 mM MES pH 5.7, 20 mM KCl, 10 mM CaCl_2_ and 0.1 % BSA for 90 min. Released mesophyll protoplasts were harvested via centrifugation at 100×*g* for 2 min, washed twice with W5 buffer (154 mM NaCl, 125 mM CaCl_2_, 5 mM KCl, 2 mM MES pH 5.7), and then resuspended in MMg buffer (0.4 M mannitol, 15 mM MgCl_2_, 4 mM MES pH 5.7) to a final concentration of 2 × 10 cells per mL.

For polyethylene glycol (PEG)-mediated transfection, 50 μg of plasmid DNA was introduced into 500 μL protoplast suspension mixed with an equal volume of PEG solution (40 % PEG4000, 200 mM mannitol, 100 mM CaCl_2_), and the mixture was incubated for 15 min. The transfection reaction was terminated by one wash with W5 buffer. After 12-16 h of incubation in W5 buffer, transfected protoplasts were centrifugation at 100×*g* for 5 min.

Total protein was extracted from pelleted protoplasts using 400 μL protein extraction buffer with vortexing. A 40-μL aliquot was reserved as the input control, while the remaining lysate was incubated with 20 μL anti-FLAG M2 affinity gel (Sigma) for 90 min to capture interacting proteins. The affinity beads were washed four times with wash buffer (10 % glycerol, 25 mM Tris-HCl pH 7.5, 1 mM EDTA, 150 mM NaCl, 0.5 % IGEPAL (Sigma-Aldrich, CA-630), and bound proteins were eluted by heating the beads in SDS loading buffer at 90 °C.

Subsequently, protein samples were resolved by SDS-PAGE and subjected to immunoblot analysis with the following antibodies: anti-FLAG-HRP (Sigma), and anti-HA-HRP (Roche).

### Phylogenetic analysis

BLASTP (BLAST 2.9.0+, e-value 10) was used to identify putative GORK orthologues within the order of the Brassicales. Subsequently, an alignment was created using the online version of MAFFT 7 (Katoh et al., 2002) and a phylogenetic analysis was performed on the CIPRES web portal using RAXML-HPC2 on XSEDE (v8.2.12) with the automatic protein model assignment algorithm using ML criterion and 250 bootstrap replicates (M. A. Miller, 2010; Stamatakis, 2014). The JTT likelihood protein model was selected as the best scoring model for ML analysis. The resulting tree was rooted using Arabidopsis SKOR, visualized using MEGA10, and edited in Corel-DRAW Home & Student x7.

### Statistical analysis

Data for quantification analyses are presented as mean ± SEM or ± SD. All analyses were performed using GraphPad Prism 9.0. The two-tailed Student’s *t*-test and one-way ANOVA followed by multiple comparisons was used for two-group comparisons and multiple comparisons.

## Acknowledgements

We thank Manuel Nieves-Cordones and Anne-Aliénor Véry for providing *gork1-2* (GABI_865F05) and *gork-2* (SALK_144737) plant materials. We acknowledge all past and current members of the Zipfel group, especially to Keran Zhai, Yukihisa Goto and Álvaro D. Fernández-Fernández, for fruitful discussions. This work was supported by the University of Zurich (C.Z.), the Gatsby Charitable Foundation (C.Z.), the European Research Council under the European Union’s Horizon 2020 research and innovation programme no. 773153 (project ‘IMMUNO-PEPTALK’) (C.Z.) and the Swiss National Science Foundation grants no. 31003A_182625, 310030_212382 and 320030_228294 (C.Z.). G.K. was supported by an SNSF Postdoctoral Fellowship (TMPFP3_224980), T.A.D. was supported by postdoctoral fellowships from the European Molecular Biology Organization (LTF-100-2017) and the Natural Sciences and Engineering Council of Canada (PDF-532561-2019).

## Author Contributions

L.W., L.Y. and C.Z conceived this study and designed the experiment. L.W. and L.Y. performed NMT experiment, split-luciferase assay and stomatal measurement and analyzed the data. R.T. and T.A.D. identified candidate BIK1 target sites. R.T. performed *In vitro* radioactive kinase assays and split-luciferase assay. S.C. conducted the co-IP experiment in protoplast and helped with plasmid construction. S.Z performed the split-ubiquitin yeast two-hybrid assay. L.W. and T.M performed TEVC experiments. G.K assisted with structure analysis and AlphaFold3 predictions and S.S performed the comparative genomics and phylogenetic analysis. L.Y., K.W., and Y.L helped with CRISPR mutant and complemented line generation. D.G and R.H hosted and assisted in the TEVC experiments. C.Z obtained funding and supervised the project. L.Y., L.W., R.T., and C.Z wrote the manuscript, with comments from all authors.

## Data availability

All data supporting the findings of this study are available within the manuscript.

---

**Figure S1.**
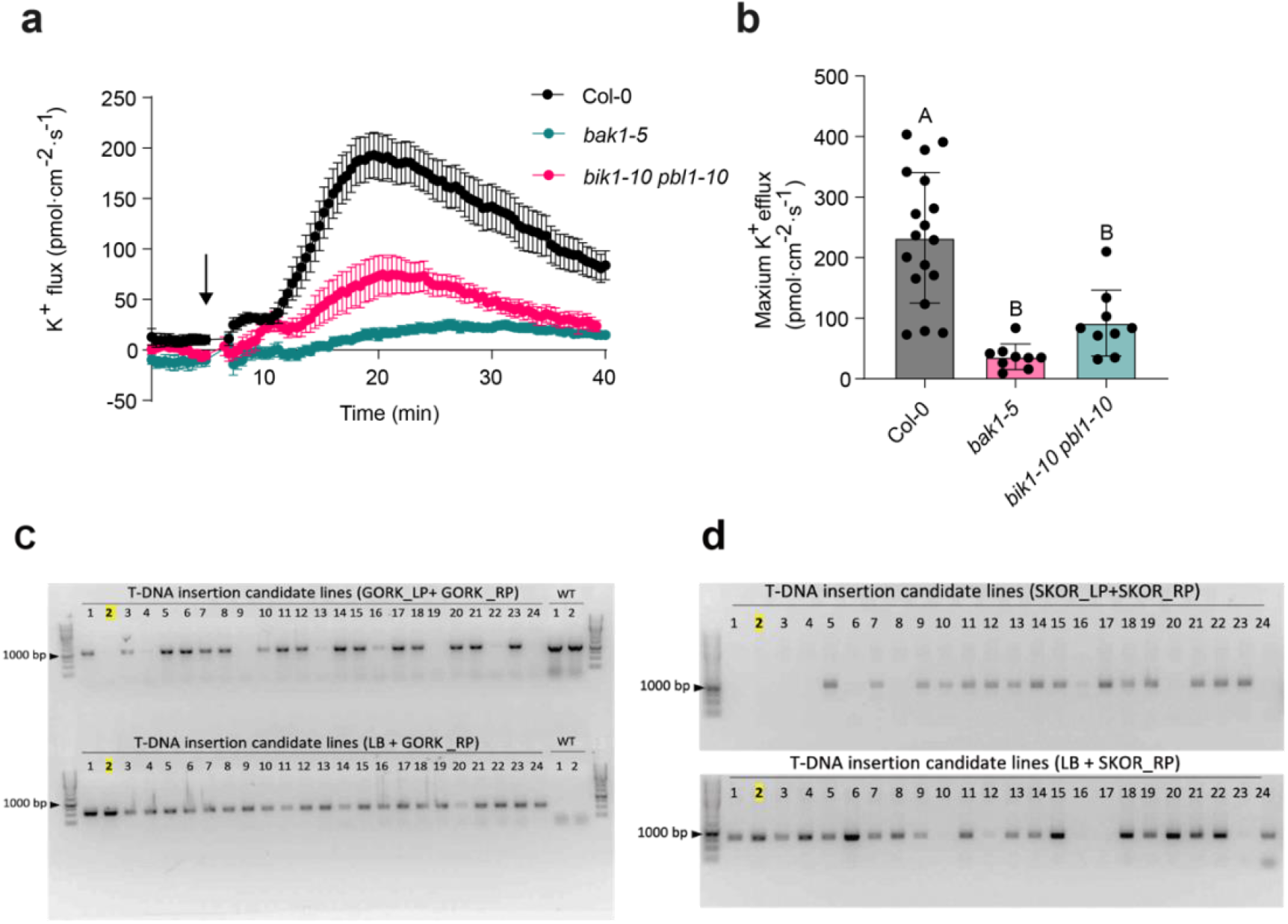
flg22-induced K^+^ efflux is BAK1- and BIK1/PBL1-dependent. **a)** Time-course recordings of K^+^ flux in mesophyll cells of Col-0, *bak1-5* and *bik1-10 pbl1-10* using NMT. Arrows indicate the addition of flg22 to the final concentration at 100 nM. Data are means ± SE, n = 9-18. **b)** The individual data point of maximum K^+^ efflux induced by 100 nM flg22 from each cell was plotted. Statistical significance was determined using one-way ANOVA followed by multiple comparisons. Different letters indicate statistical differences. **c-d)** Genotyping PCR identification of *gork-2 skor-2* double T-DNA insertion mutant. The lane highlighted in yellow marks the homozygous line selected for subsequent experiments. **c)** PCR genotyping for candidate lines of *gork-2*. Upper panel: PCR using gene-specific primers GORK_LP + GORK_RP; Lower panel: PCR using T-DNA border primer LB paired with gene-specific reverse primer GORK_RP, WT served as control. **d)** PCR genotyping for candidate lines of *skor-2*. Upper panel: PCR using gene-specific primers SKOR_LP + SKOR_RP; Lower panel: PCR using T-DNA primer LB together with SKOR_RP; Lanes 1-24 represent independent T-DNA insertion candidate seedlings.

**Figure S2.**
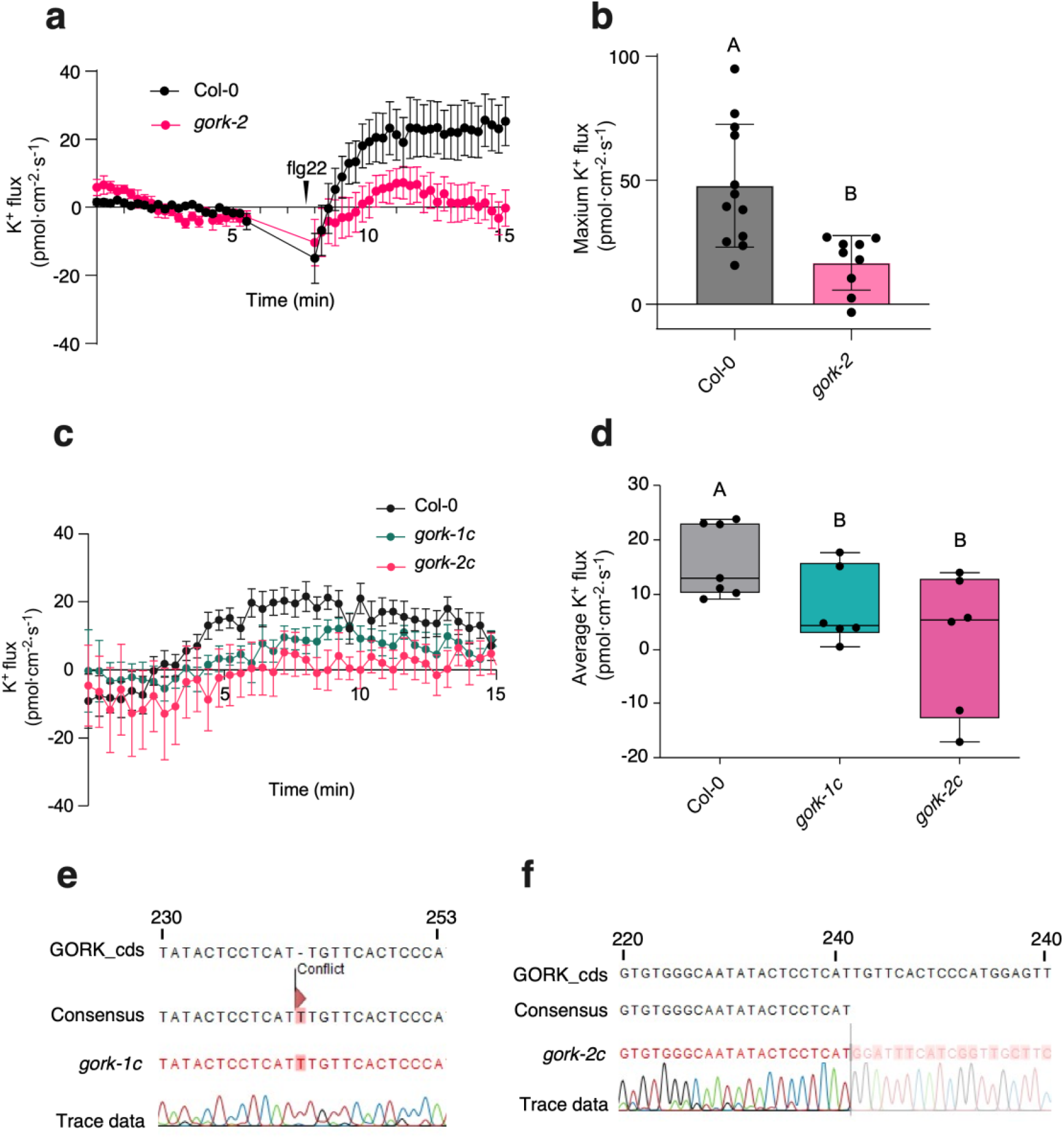
flg22-triggered K^+^ efflux is reduced in guard cells of *gork* mutants. **a)** K⁺ flux measurements in guard cells of Col-0 and *gork-2* plants. Positive values indicate K⁺ efflux. Arrows indicate the addition of flg22 to a final concentration of 100 nM. Data are presented as means ± SE. **b)** Quantification of the maximum K⁺ efflux shown in (a). Data are presented as means ± SE (n = 9-12 guard cells). Statistical significance was performed using Student’s *t*-test. Different letters indicate statistically significant differences (*p* < 0.05). **c)** K⁺ flux measurements in guard cells of the indicated genotypes. K⁺ flux recording was initiated 1 min after treatment with 100 nM flg22. Data are presented as means ± SE. **d)** Quantification of the average K⁺ flux during 5-15 min after flg22 treatment from (c). Experiments were repeated twice with 6-7 individual plant from each genotype. Data are presented as means ± SE. Statistical significance was determined using one-way ANOVA followed by multiple comparisons. **e)** Genotyping of the CRISPR-generated *gork-1c* mutant. A single nucleotide insertion (T) after position 241 bp causes a frameshift and premature stop codon. **f)** Genotyping of the CRISPR-generated *gork-2c* mutant. A 196-bp deletion spanning positions 242-437 bp causes a frameshift and premature stop codon.

**Figure S3.**
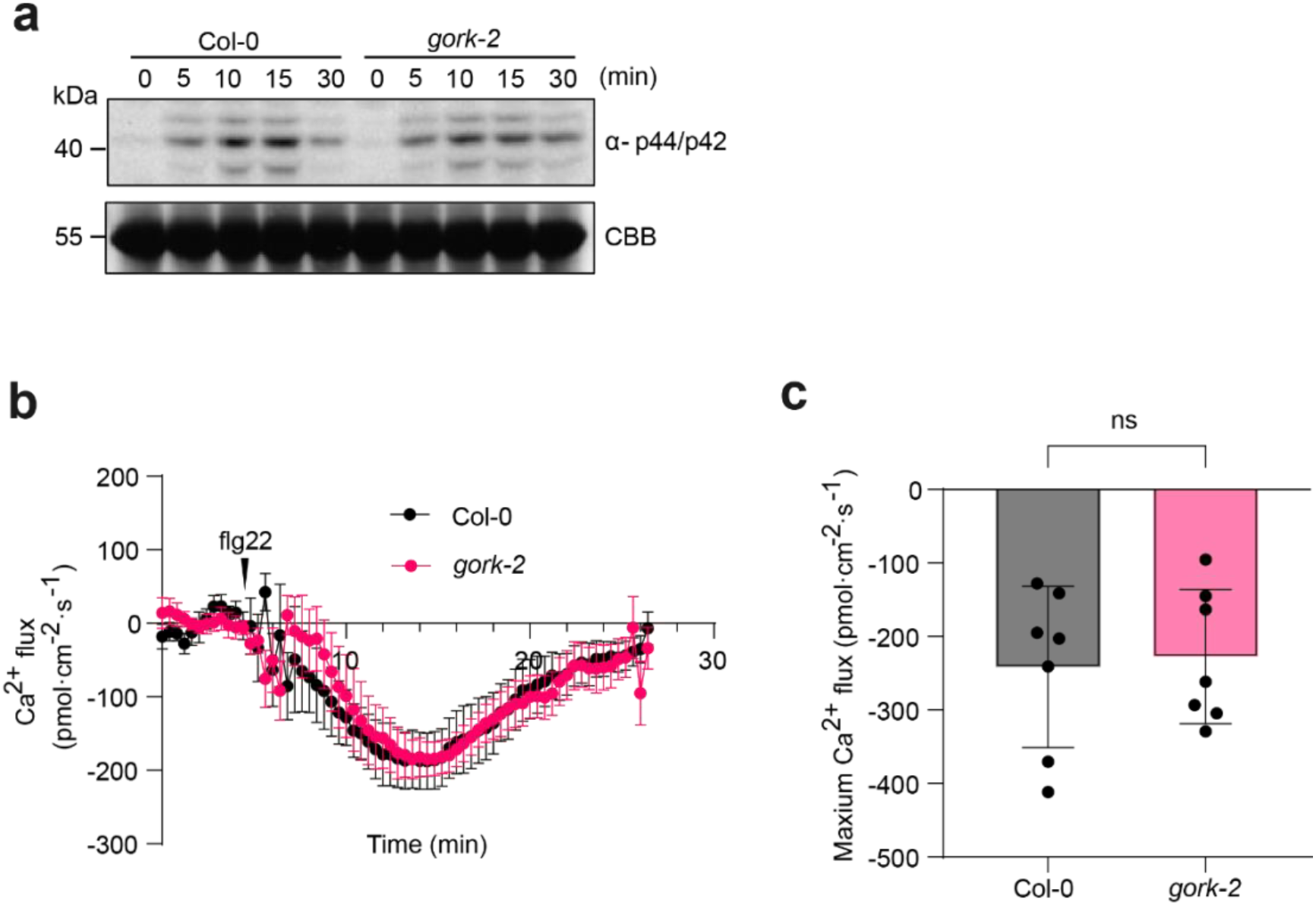
flg22-induced MAPK activation and Ca^2+^ elevation are not affected in *gork-2* mutant. **a)** MAPK activation induced by 100 nM flg22 at the indicated time points in Col-0 and *gork-2* plants. CBB staining was used as a loading control. Experiments were independently repeated twice with similar results. **b)** Time-course analysis of flg22-induced Ca^2+^ flux in mesophyll cells. Arrows indicate the addition of flg22 to a final concentration of 100 nM. Data are presented as means ± SE. **c)** Quantification of the maximum Ca^2+^ flux shown in (b). Experiments were independently repeated twice, with seven individual leaves from each genotype analyzed. Data are presented as means ± SE. Statistical significance was determined using Student’s *t*-test.

**Figure S4.**
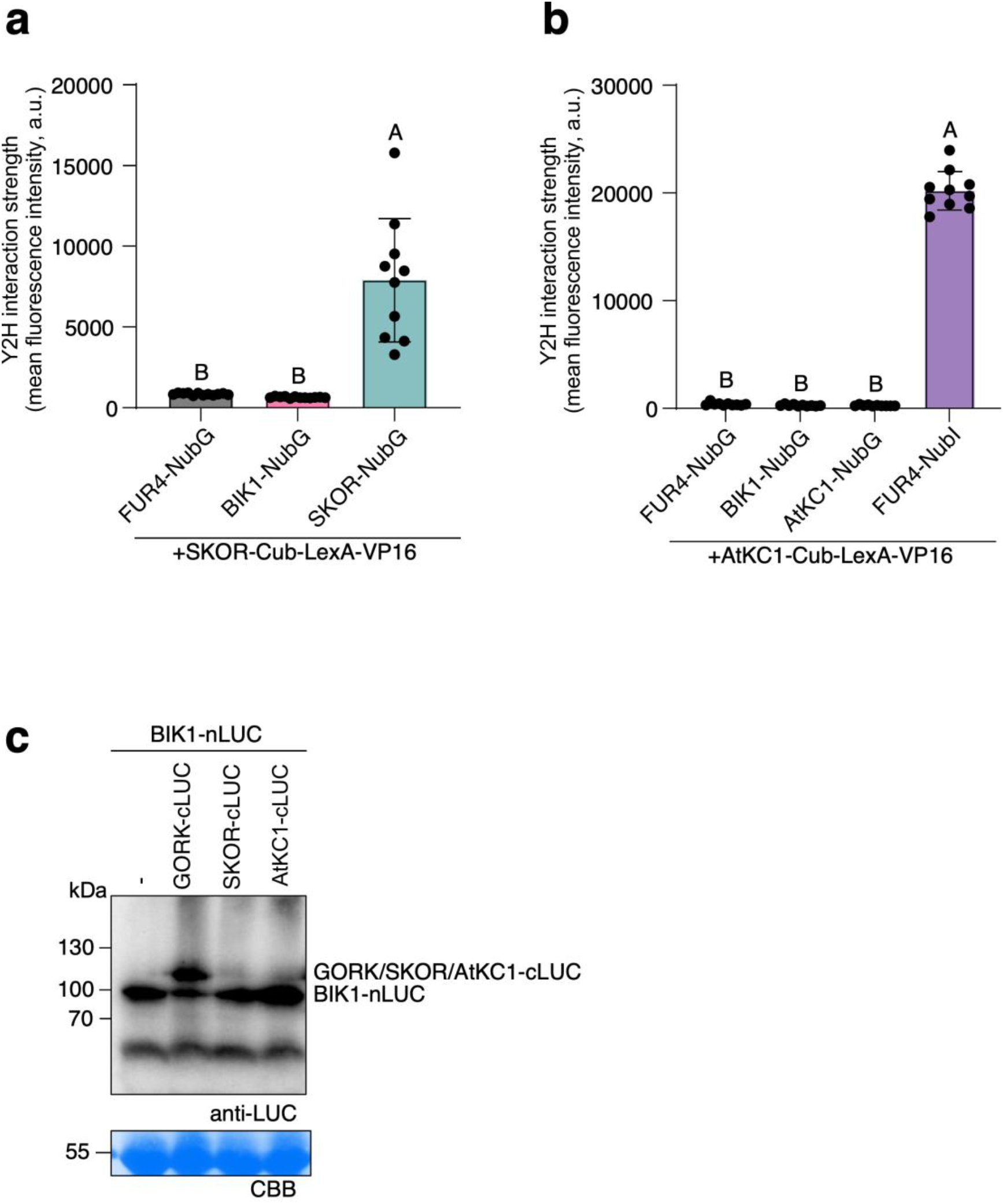
SKOR does not interact with BIK1. **a-b)** Split-ubiquitin yeast two-hybrid assay between indicated proteins. FUR4-NubG was used as negative control. **a)** Yeast cells co-expressing SKOR-Cub-LexA-VP16 together with the indicated NubG-fused membrane proteins were subjected to flow cytometry analysis. n = 10-12, error bars represent ± SE. **b)** Yeast cells co-expressing AtKC1-Cub-LexA-VP16 together with the indicated NubG-fused membrane proteins were subjected to flow cytometry analysis. n = 10, error bars represent ± SE. For a-b statistical significance was determined using one-way ANOVA with multiple comparisons. Different letters indicate statistical differences. **c)** Anti-LUC western blot indicating expression of GORK, SKOR and AtKC1-cLUC tagged variants along with BIK1-nLUC.

**Figure S5.**
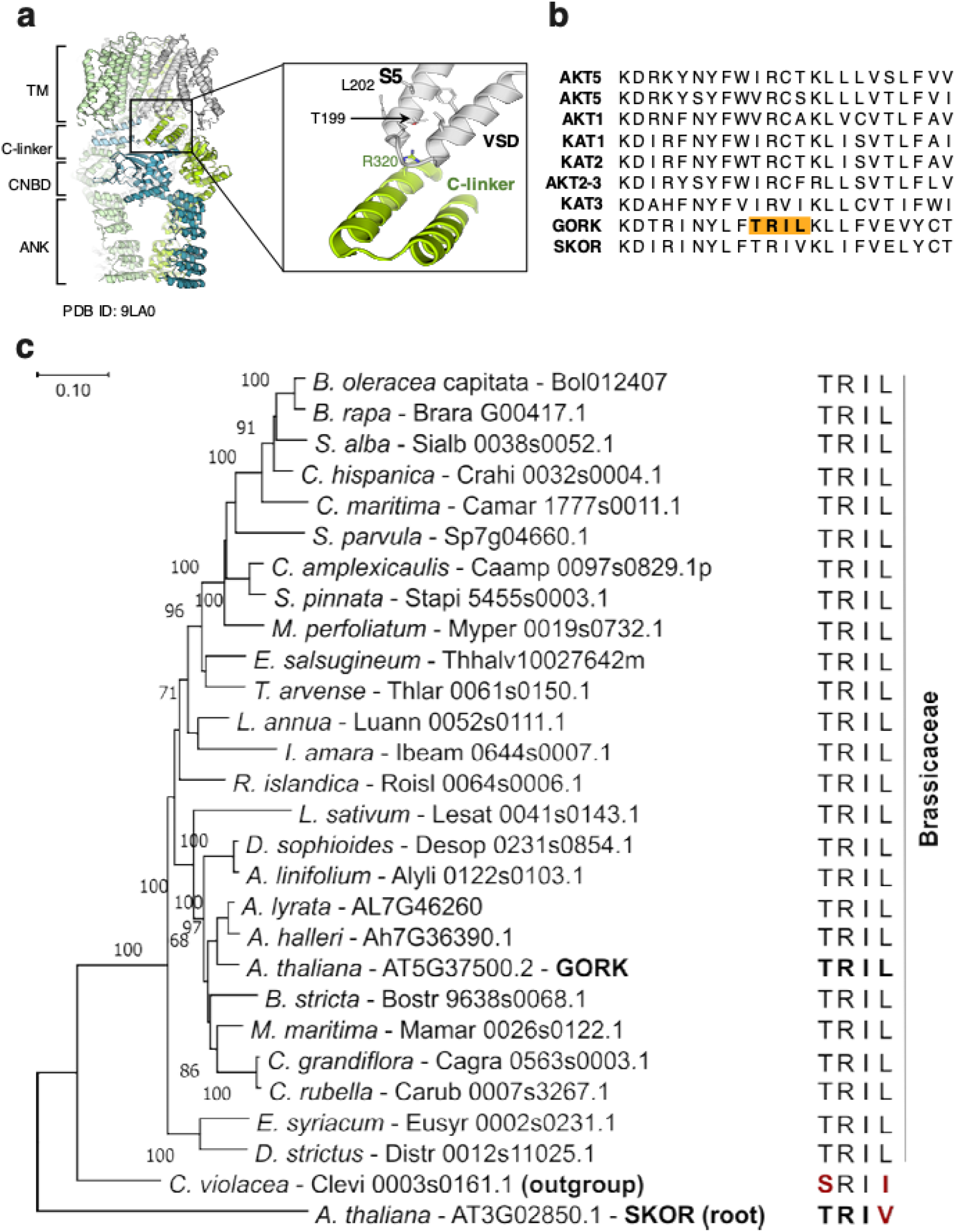
Structural localization and evolutionary conservation of T199. **a)** Structural representation of Arabidopsis GORK (PDB ID: 9LA0). Domains are annotated: TM, transmembrane region; C-linker; CNBD, cyclic nucleotide-binding domain; ANK, ankyrin repeat domain. The enlarged view highlights the VSD (voltage-sensing domain) and C-linker region, indicating the position of conserved T199 residue **b)** Multiple sequence alignment of plant Shaker K^+^ channels, revealing that the T199 residue (within the TRIL motif) is highly conserved among GORK clade outward K^+^ channels. **c)** A phylogenetic analysis of putative GORK orthologues in the family of the Brassicaceae and *C. violaceae* (order of the Brassicales). Maximum-likelihood analysis bootstrap values are indicated, and only values higher than 65 are shown. The scale bar represents 0.1 AA substitutions per site. Arabidopsis SKOR was incorporated in the analysis and used to root the phylogenetic tree. The conserved sequence motif corresponding to the T199-containing region is displayed on the right of each protein. *Arabidopsis thaliana* SKOR and *C. violacea* Clevi 0003s0161.1 were used as outgroups.

**Figure S6.**
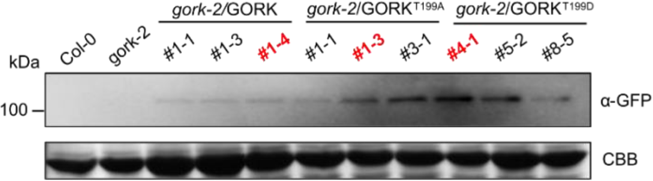
Protein accumulation in transgenic GORK complementation lines. CBB staining serves as the total protein loading control. Lines marked with red were selected for subsequent experimental analyses. Col-0: wild-type Arabidopsis; *gork-2*; *gork-2*/GORK, *gork-2*/GORK^T199A^, and *gork-2*/GORK^T199D^ represent *gork-2* mutant plants transformed with wild-type GORK, GORK^T199A^ and GORK^T199D^ -GFP constructs driven by the native *GORK* promoter, respectively.

**Figure S7.**
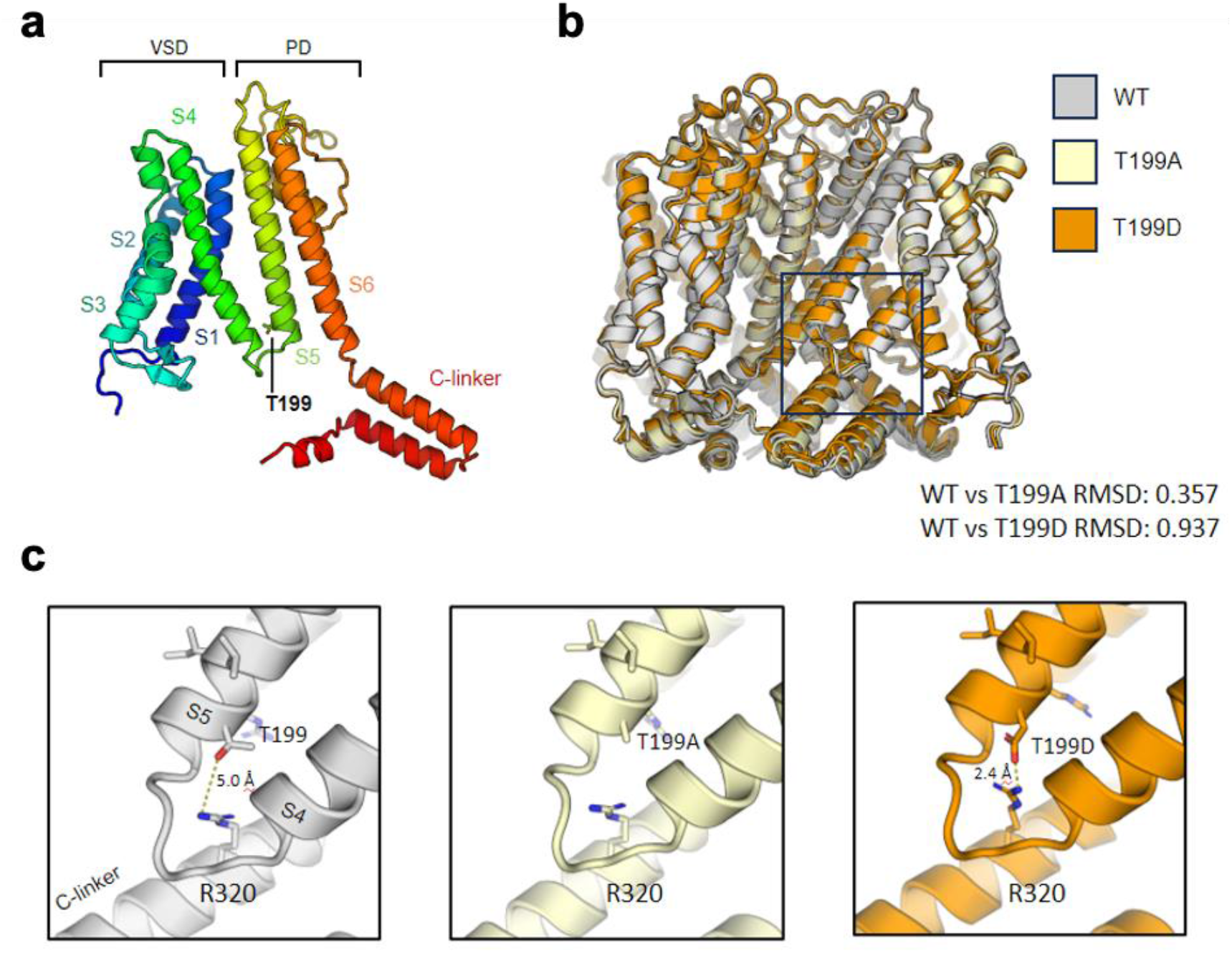
Structural comparison of GORK, GORK^T199D^ and GORK^T199A^. **a)** Domain organization of a monomer and localization of mutation site T199. **b)** The predicted structures were structurally aligned to the GORK WT model, and the root-mean-square deviation (RMSD) was shown to indicate structural similarity. **c)** Close-up views of the T199-containing S4–S5 linker region and its surrounding residues in the GORK WT, GORK^T199D^ and GORK^T199A^. Native threonine T199 only forms weak long-range contacts with R320. Alanine substitution (T199A) removes the polar side chain and disrupts native intermolecular contacts. Aspartate substitution (T199D) introduces a negatively charged carboxylate group, which generates tight electrostatic attraction with the positively charged R320, narrowing the separation between two helices. All structures were predicted using AlphaFold3 (AF3) and visualized using PyMOL.

