## Supplementary Tables S1-S3 for "The *Arabidopsis* outward K^+^ channel GORK mediates flg22-induced K^+^ efflux and stomatal closure"

**Supplementary Table S1. Genotypes used in this study**

| **Genotype** | **Details** | **Origin/Reference** |
| --- | --- | --- |
| *gork-2* | T-DNA insertion mutant of GORK, SALK_144737 | SALK collection |
| *skor-2* | T-DNA insertion mutant of GORK, GABI_391G12 | (Nieves-Cordones et al., 2019) |
| *gork-2 skor-2* | GORK and SKOR double mutant generated by crossing gork-2 and skor-2 | This study |
| *bik1-10 pbl1-10* | CRISPR/Cas9-generated BIK1 PBL1 double mutant | (Song et al., 2026) |
| *bak1-5* | EMS-induced C408Y missense mutation | (Schwessinger et al., 2011) |
| *gork-1c* | CRISPR/Cas9-generated GORK knockout line | This study |
| *gork-2c* | CRISPR/Cas9-generated GORK knockout line | This study |
| *gork-2*/GORK | gork-2 complemented with *pGORK::GORK-GFP* | This study |
| *gork-2*/GORK^T199A^ | gork-2 complemented with *pGORK::GORK*^T199A^*-GFP* | This study |
| *gork-2*/GORK^T199D^ | gork-2 complemented with *pGORK::GORK*^T199D^*-GFP* | This study |

**Supplementary Table S2. Constructs used in this study**

| **Construct** | **Usage** | **Cloning method** | **Destination** | **Origin/Reference** |
| --- | --- | --- | --- | --- |
| *FUR4-NubG* | Prey in split-ubiquitin assay | Golden gate cloning | CZLp9747 | This study |
| *GORK-NubG* | Prey in split-ubiquitin assay | Golden gate cloning |  | This study |
| *SKOR-NubG* | Prey in split-ubiquitin assay | Golden gate cloning |  | This study |
| *AtKC1-NubG* | Prey in split-ubiquitin assay | Golden gate cloning |  | This study |
| *BIK1-NubG* | Prey in split-ubiquitin assay | Golden gate cloning |  | This study |
| *BIK1-Cub-LexA-VP16* | Bait in split-ubiquitin assay | Golden gate cloning | CZLp9748 | This study |
| *GORK -Cub-LexA-VP16* | Bait in split-ubiquitin assay | Golden gate cloning |  | This study |
| *GORK-cLUC* | Split-luciferase complementation assay | Golden gate cloning | pICSL86955OD | This study |
| *SKOR-cLUC* | Split-luciferase complementation assay | Golden gate cloning | pICSL86955OD | This study |
| *AtKC1-cLUC* | Split-luciferase complementation assay | Golden gate cloning | pICSL86955OD | This study |
| *BIK1-nLUC* | Split-luciferase complementation assay | Golden gate cloning | pICSL86955OD | This study |
| *BIK1-HA* | Transient expression in Arabidopsis protoplast | Golden gate cloning | pICSL50009 | This study |
| *ATKC1-FLAG* | Transient expression in Arabidopsis protoplast | Golden gate cloning | pICSL50007 | This study |
| *GORK-FLAG* | Transient expression in Arabidopsis protoplast | Golden gate cloning | pICSL50007 | This study |
| *MBP-BIK1* | *In vitro* kinase assays | In fusion cloning | pOPINM | This study |
| *MBP-BIK1** | *In vitro* kinase assays | In fusion cloning | pOPINM | (Toth et al., 2026) |
| *MBP-GORK* | *In vitro* kinase assays | In fusion cloning | pOPINM | This study |
| *MBP-SKOR* | *In vitro* kinase assays | In fusion cloning | pOPINM | This study |
| *MBP-GORK^T199A^* | *In vitro* kinase assays | In fusion cloning | pOPINM | This study |
| *GORK-YFP* | Heterologous expression in Xenopus oocytes | Restriction digestion and ligation | pGEMHE | This study |
| *GORK^T199A^-YFP* | Heterologous expression in Xenopus oocytes | Restriction digestion and ligation | pGEMHE | This study |
| *GORK^T199D^-YFP* | Heterologous expression in Xenopus oocytes | Restriction digestion and ligation | pGEMHE | This study |
| *BIK1-YFP* | Heterologous expression in Xenopus oocytes | Restriction digestion and ligation | pGEMHE | This study |
| *BIK1*-YFP* | Heterologous expression in Xenopus oocytes | Restriction digestion and ligation | pGEMHE | This study |
| *pGORK::GORK-GFP* | Genetic complementation of ***gork-2* with native promoter** | Golden gate cloning | pICSL86955OD | This study |
| *pGORK::GORK^T199A^-GFP* | Genetic complementation of ***gork-2* with native promoter** | Golden gate cloning | pICSL86955OD | This study |
| *pGORK::GORK^T199D^-GFP* | Genetic complementation of ***gork-2* with native promoter** | Golden gate cloning | pICSL86955OD | This study |

**Supplementary** **Table S3. Primers used in this study**

| **Primer name** | **Sequence (5’-3’)** | **Usage** | **Origin/Reference** |
| --- | --- | --- | --- |
| GORK1_BpiI_GATC_F | atGAAGACTTgatcATGGGTAGACTCCGGAG | *Xenopus* oocytes expression | This study |
| GORK_BpiI_AATT_R | ATGAAGACTTaattTTATGTTTGATCAGTAGTATCACTGATCAT |  | This study |
| BIK1_BamHI_F | ccggggatccatgggttcttgcttcagttct |  | This study |
| BIK1_EcoRI_R | tagagaattcTTAcacaaggtgcctgccaaaag |  | This study |
| *gork*-KO-R | GCATCACCTAGAAACGAACCA | Genotyping | This study |
| *gork*-KO-F | TTGGTTCCACTAAAGCTGCC |  | This study |
| *gork*-tDNA-RP | TTTTGGTTCTGATTCGGTTTG |  | This study |
| *gork*-tDNA-LP | ACAAATGATCCACGACGTTTC |  | This study |
| *skor*-tDNA-LP | ACACGATCATTCCCATTCTTG |  | This study |
| *skor*-tDNA-RP | TATGAACCGAAACAAACTCGG |  | This study |
| LBb1.3 | ATTTTGCCGATTTCGGAAC |  | This study |
| GORK sgRNA1-F | ATTGCTCCATGGGAGTGAACAATG | CRISPR/Cas9 | This study |
| GORK sgRNA1-R | AAACCATTGTTCACTCCCATGGAG |  | This study |
| GORK sgRNA2-F | ATTGCCTTATAAATAAGATCCCAA |  | This study |
| GORK sgRNA2-R | AAACTTGGGATCTTATTTATAAGG |  | This study |
| GORK_T-A-sense | CTATTCGCTAGAATCTTAAAGCTCTTGTTCGTTGAAGTTTATTGT | Mutagenesis | This study |
| GORK_T-A-antisense | GATTCTAGCGAATAGATAGTTGATTCTTGTGTCTTTCTC |  | This study |
| GORK-T-D-sense | CTATTCGATAGAATCTTAAAGCTCTTGTTCGTTGAAGTTTATTGT |  | This study |
| GORK-T-D-antisense | GATTCTatcGAATAGATAGTTGATTCTTGTGTCTTTCTC |  | This study |
| BIK1_C_F | ATCGTCTCTTACGGGTCTCACACCATGGGTTCTTGCTTCAGTTC | Split-ubiquitin yeast two-hybrid | This study |
| BIK1_D_R | ATCGTCTCTCAGAGGTCTCACCTTCACAAGGTGCCTGCC |  | This study |
| GORK_C_F | ATCGTCTCTTACGGGTCTCACACCATGGGTAGACTCCGGAG |  | This study |
| GORK_D_R | ATCGTCTCTCAGAGGTCTCACCTTTGTTTGATCAGTAGTATCACTGATC |  | This study |
| SKOR_C_F | ATCGTCTCTTACGGGTCTCACACCATGGGAGGTAGTAGCGG |  | This study |
| SKOR_D_R | ATCGTCTCTCAGAGGTCTCACCTTTGTTTCAACAGCCAAATACAG |  | This study |
| ATKC1_C_F | ATCGTCTCTTACGGGTCTCACACCATGTCTACGACGACTACTGAG |  | This study |
| ATKC1_D_R | ATCGTCTCTCAGAGGTCTCACCTTGAAAATATATAAATGATCGTTTTC |  | This study |
| Fur4p_C_F | ATCGTCTCTTACGGGTCTCACACCatgccagacaatctatcattac |  | This study |
| Fur4p_D_R | ATCGTCTCTCAGAGGTCTCACCTTgccaatgaaagtcttttcgtg |  | This study |
| Fur4p_mut_F | ATCGTCTCTgacAtttgcgccaaataatgtg |  | This study |
| Fur4p_mut_R | ATCGTCTCTTgtctcattggaagattccataag |  | This study |
| BIK1-F | ATCGTCTCTTACGGGTCTCAAATGGGTTCTTGCTTCAGTTCTC | Protoplast transient expression | This study |
| BIK1-R | ATCGTCTCTCAGAGGTCTCACGAAccCACAAGGTGCCTGCCAAAAG |  | This study |
| GORK_pOPINM_F | AAGTTCTGTTTCAGGGCCCGTTTCGGGTTCGCAAAGTGGT | Recombinant Proteins | This study |
| GORK_pOPINM_R | ATGGTCTAGAAAGCTTTATAAGATTCTAGTGAATAGATA |  | This study |
| SKOR_pOPINM_F | AAGTTCTGTTTCAGGGCCCGTATCGAGTTCATAGAGTAAT |  | This study |
| SKOR_pOPINM_R | ATGGTCTAGAAAGCTTTAGACGATTCTTGTAAAAAGGT |  | This study |
